# Diversified Portfolios of Bacterial Communities in Peri-urban Miyawaki Forests and Grassland Shaping Ecosystem Functions and Services

**DOI:** 10.64898/2026.02.10.704982

**Authors:** Kanika Bansal, Indira Singh, V. Vaishnavi, Reddy R.L. Raghunath, Aadya Joshi, Gayatri Bakhale, Jagdish Krishnaswamy

## Abstract

Soil microflora is fundamental to ecosystem functioning, yet their contribution in Miyawaki afforestation, a globally implemented ecological engineering approach, remains poorly characterized. In the present study, we examined the bacterial taxonomic diversity and their functional potential in a peri-urban Miyawaki mini-forest and compared with a nearby grassland the pre-existing ecosystem across dry and wet seasons. The Miyawaki plantation comprised of highly diverse native trees, sub-trees and shrubs spanning evergreen and deciduous varieties, potentiating nitrogen-fixation, diverse litter generation and rooting strategies resulting in pronounced functional heterogeneity. Notably relative to grassland, the Miyawaki forest was intensively managed and supplemented with organic amendments, and supportive irrigation, buffering the seasonal moisture stress. Using 16S rRNA amplicon sequencing of soil eDNA, we characterized seasonal variation in soil bacterial communities in both the systems. The observed soil bacterial community organization in forest as compared to the unmanaged grassland indicates combined influence of vegetation structure, dense canopy cover, continuous litter generation and root exudates. Microbial assemblages in the forest specialised in heterotrophic complex carbon degradation, biofilm formation, exopolysaccharide production and sporulation pathways which suggests adaptive abilities to anoxic microsites and other stressful conditions. In contrast, grassland soils harboured less diversified bacterial communities dominated by phototrophic and oxidative stress adaptation pathways consistent with sun lit, non-irrigated and moisture-variable conditions. Nonetheless, functional divergence in dry season reflects temporal reorganization of microbial communities marking a gradual trend towards soil ecosystem development. Together, these findings establish microbial baseline for Miyawaki forests revealing how tree-dense mini-forests restructure soil bacterial communities relative to grasslands highlighting the value of identifying soil microbial indicators for critically evaluating urban afforestation outcomes over extended time scales to inform sustainable design and policy.

**Graphical abstract:** 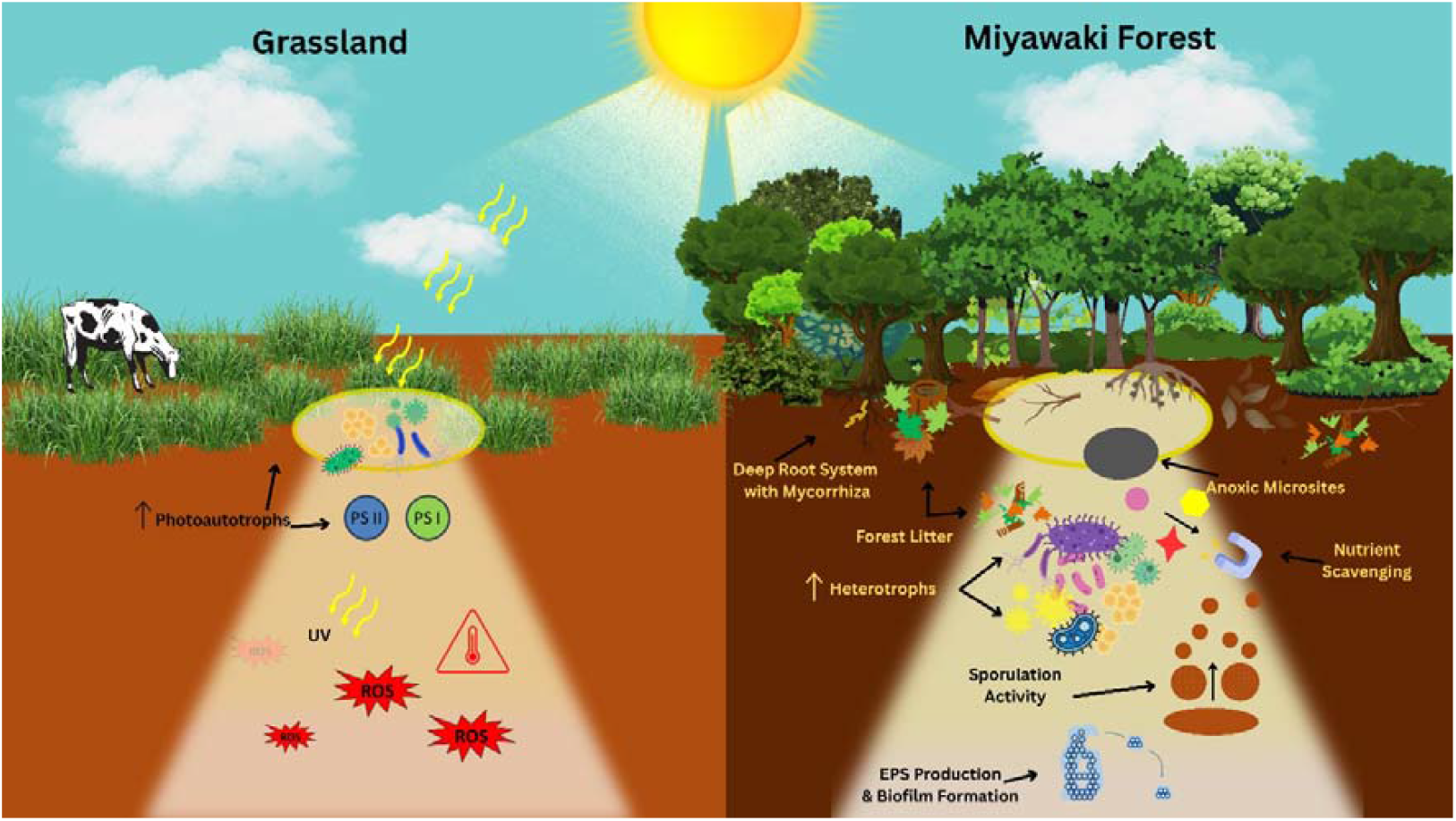

Schematic representation of the study depicting the Miyawaki forest and nearby grassland.

## Introduction

Worldwide deforestation has reached an alarming rate of 10 million hectares per year (FAO and UNEP, 2020). This escalates the risk of natural disasters and poses serious consequences to the local as well as the broader regional ecology. According to the Intergovernmental Panel on Climate Change (IPCC) to augment the ecosystem sustainability, efficient restoration, afforestation and reforestation efforts are the most practiced options globally (Intergovernmental Panel On Climate Change (IPCC), 2023). However, traditional techniques of reforestation might need hundreds of years to reestablish the ecosystems (Di Sacco et al., 2021). In the current scenario, with high rates of environmental degradation, effective human interventions are needed to accelerate and establish mature ecological systems. The Miyawaki method proposed by the Japanese botanist Professor Akira Miyawaki in the 1970s, is an active ecological engineering approach to create diverse, compact and self-sustaining native forests (Miyawaki, 1999). Compared to decades needed for extensive forest succession, with Miyawaki method the climax ecosystem can be reached within 15-20 years of onset (Lewis, 2022). Essentially, the Miyawaki method is based on the principle of ‘native forests by native trees’ i.e., reviving the native vegetation present in the ecosystem prior to human interventions. The method relies on dense plantation, intensive soil preparation, organic amendments and sustained irrigation during establishment accelerating biomass accumulation and canopy closure to rapidly grow the forest-like structure. Though, they differ fundamentally from natural forest succession, where ecosystem structure and function emerge from long-term biotic and abiotic feedback. Till date, more than 3000 Miyawaki primary forests have been developed containing over 40 million native trees worldwide (Vaidya, 2025). These mini “forests” are portrayed as an oasis of biodiversity and potential environmental healers, that also support reconnecting the local communities with nature. Yet, critical assessment of their ecological effectiveness, biodiversity benefits and resilience is required, as questions remain about site suitability, resource intensity, ecological mismatches and long-term outcomes of dense plantations versus natural forests (Morales et al., 2025; Tushar Kaushik, 2019; Vaidya, 2025).

Microbial communities are the key drivers for ecological successions, performing indispensable roles in nutrient cycling, soil structuring and plant-microbe interactions leading to successful reforestation (Fierer & Jackson, 2006). Across reforestation, afforestation or urban greening initiatives, vegetation is the dominant driver of soil microbial community and its function depicted by litter inputs, root exudation, soil moisture and microclimatic modification (Wardle et al., 2004). Comprehensive microbial community assessment of largest planted forest in Saihanba region implied complex network of microbial community as compared to the grassland soil, highlighting a tight coupling of microbial community with the vegetation types (K.-C. Huang et al., 2024). A meta-analysis across reforestation studies showed that afforestation often increases soil enzymatic potential influencing microbial functions strongly than their diversity (H. Huang et al., 2022). Chronosequence and long-term studies indicate that microbial communities undergo successional turnover over years and decades with an earlier stabilization of functional attributes relative to taxonomic composition, likely driven by a need for functional redundancy and environmental filtering (Fierer et al., 2010; K.-C. Huang et al., 2024; Jurburg et al., 2017). Especially in early-stage forests under afforestation models like Miyawaki, microbes act as ecosystem engineers. Bacterial metabolic activities like biofilm formation have been directly linked to rhizosphere chemistry by buffering microclimatic fluctuations and sequestering heavy metals (Barra Caracciolo & Terenzi, 2021; Keiluweit et al., 2018). Specifically, bacterial communities not only influence soil health, but also lay the biochemical foundation for symbiotic fungal colonization, later enabling mycorrhizal networks for facilitating the mature forests.

Advancements in high-throughput sequencing and metagenomics have revolutionized our understanding of how these communities assemble in complex environments such as soil. Unlike culture-based approaches, metagenomics provides direct insights into taxonomic and functional signatures from environmental DNA (eDNA), allowing in depth exploration of ecosystems. Post landscape restorations metagenomics has proven useful, for instance during the reforestation of decommissioned cranberry bogs in New Jersey Pine Barrens, shotgun metagenomics revealed microbial successions linked to vegetation type and hydrology (Eaton et al., 2017). These are indicators of enhanced critical guilds of soil biota and increased nutrient availability in the restored habitats. Hence, interactions between plants, soil and microorganisms are the major drivers of functions of forest ecosystems and modifications in plant cover tremendously shift the soil microbial compositions.

Previous studies on Miyawaki afforestation have majorly focused on vegetation, protocol used for set-up and ecosystem services outcomes. For instance, the importance of soil preparation with compost and mycorrhizal inoculates coupled with high-density native plantations result in denser Miyawaki forests as compared to natural ones (Parikh & Nazrana, 2023; Swapna, 2023). High tree densities of diverse tree communities increases the competition for light, often resulting in rapid increase in plant height (Swapna, 2023). Forest establishment consistently relies on dense plantations of site-specific native tree species representing multiple forest strata for canopy, sub-canopy, sub-trees and shrubs. Globally, Miyawaki style afforestation have been adopted in parts of Europe and Southeast Asia to deliver ecosystem services such as microclimate regulation, soil stabilization and urban green cover (Da & Song, 2008; Frattaroli et al., 2017; Kiboi et al., 2014). In India, the Miyawaki method has been applied across a range of ecological and land-use contexts, including mining landscapes in Odisha, urban settings in Rajkot, and institutional campuses in Coimbatore. These implementations predominantly employed regionally native deciduous and evergreen tree species such as: *Ficus* spp., *Terminalia* spp., *Syzygium* spp., *Azadirachta indica, Mangifera indica, Cassia fistula* etc. to mimic natural vegetation (Goveanthan et al., 2019; Lagariya & Kaneria, 2021; Ranjan et al., 2016). In addition to analysing the forest establishment,, studies have assessed socio-economic aspect of the forest in terms of improved community engagement and, mental-health benefits (Rajadurai, 2024). Meta-analytic study found that plantations with mixed species tend to outperform the single species plantations by enhancing soil nutrients, microbial diversity, enzymatic activity and functional gene repertoires related to carbohydrate and polymer degradation (H. Huang et al., 2022). Mechanistically these are driven by: heterogeneity in litter quality selecting for saprophytic microbial guilds, leguminous trees improving soil nitrogen levels, diversity in root-depth impacting moisture regimes and oxygen microsites enhancing the complexity of microbial networks thus overall, nurturing a functionally diverse forest soil (Li et al., 2024).

Monitoring Miyawaki forests is typically centred around repeated assessments of vegetation structure, soil physical and chemical properties and selected faunal indicators with surveys conducted across seasons to capture the successional dynamics (C.L. Narraway et al., 2024). Yet, there are limited empirical evidences beyond the short-term vegetation metrics regarding direct implications of Miyawaki method on ecosystem services like: biodiversity recovery, temperature impacts, carbon sequestration, soil microbial functioning, nutrient cycling, moisture buffering and plant-microbe feedbacks (Morales et al., 2025). These frameworks predominantly rely on plant-centric and bulk soil indicators for ecosystem health, while studies involving a comprehensive microbial profiling and analysis of community shifts in soil obtained from Miyawaki forests are limited. However, a preliminary report suggests that in addition to the improvement in soil parameters and nutrients, Miyawaki forest soil was found to be enriched in *Paenibacillus jamilae* and *Paenibacillus polymyxa* (Deepalakshmi & Umadevi, 2023). In addition to this, Guo et al, have enumerated enrichment of culturable bacterial and fungal communities in Miyawaki plots, underscoring their importance as plant growth promoters (Guo et al., 2018). Hence, despite their critical relevance in early successional soil development and long-term restorations, details on bacterial community structure, diversity or functional guilds largely remains unexplored. This is a critical knowledge gap, as these urban afforestation efforts lead to accelerated biomass accumulation and high species density resulting in unique microclimatic conditions that also shape soil microbial consortia. In the present study, we have evaluated the ability of Miyawaki to foster early-stage recovery of soil microbial diversity and their functional composition as compared to the neighbouring grassland as reference across wet and dry seasons. The study was conducted at a 10-acre warehouse site in Mohadi, Nashik, of which 1.5-acre was converted into Miyawaki forest. Prior to the forest establishment, the whole warehouse had been under grape cultivation and subsequently left uncultivated for approximately nine years (2013-2022). In 2022, the Miyawaki plantation was initiated, while the adjacent area which had soon developed into a grassland was retained as such. This allowed a direct comparison with the surrounding unplanted grassland representing the pre-existing baseline, thus addressing the critical limitation noted among previous Miyawaki forest evaluations (Morales et al., 2025). Since, soil microbial communities and their functional potential respond rapidly to vegetation driven shifts in moisture, redox conditions, and substrate quality, they form early response indicators of ecosystem change (Bardgett & Van Der Putten, 2014). Hence, we specifically investigated tree-driven changes in litter inputs, microclimate and soil structure, which are directly reflected in bacterial community assemblages across seasons.

## Results

### Spatiotemporal vegetation trajectories before and after Miyawaki establishment

Temporal remote sensing revealed distinct land-cover trajectories between Miyawaki restoration site and adjacent grassland over the period of 10 years (2014-2024) (Figure 1(A)). Prior to the afforestation, both sites were used for grape cultivation (2014), followed by an extended untended grassland phase. Following the initiation of Miyawaki plantation in May 2022, the restored plot exhibited a rapid and visibly discernible increase in vegetation density, culminating in a canopy cover by May 2024. In contrast, the adjacent grassland retained open, sparsely vegetated structure with minimal change across the same timeframe. This sharp divergence in vegetation structure between two sites post-2022 provides clear spatial evidence of above-ground biomass accumulation in the Miyawaki forest site, while the grassland was stable, indicating comparable legacy conditions, hence it was considered as a reference point in the study.

**Figure 1.**
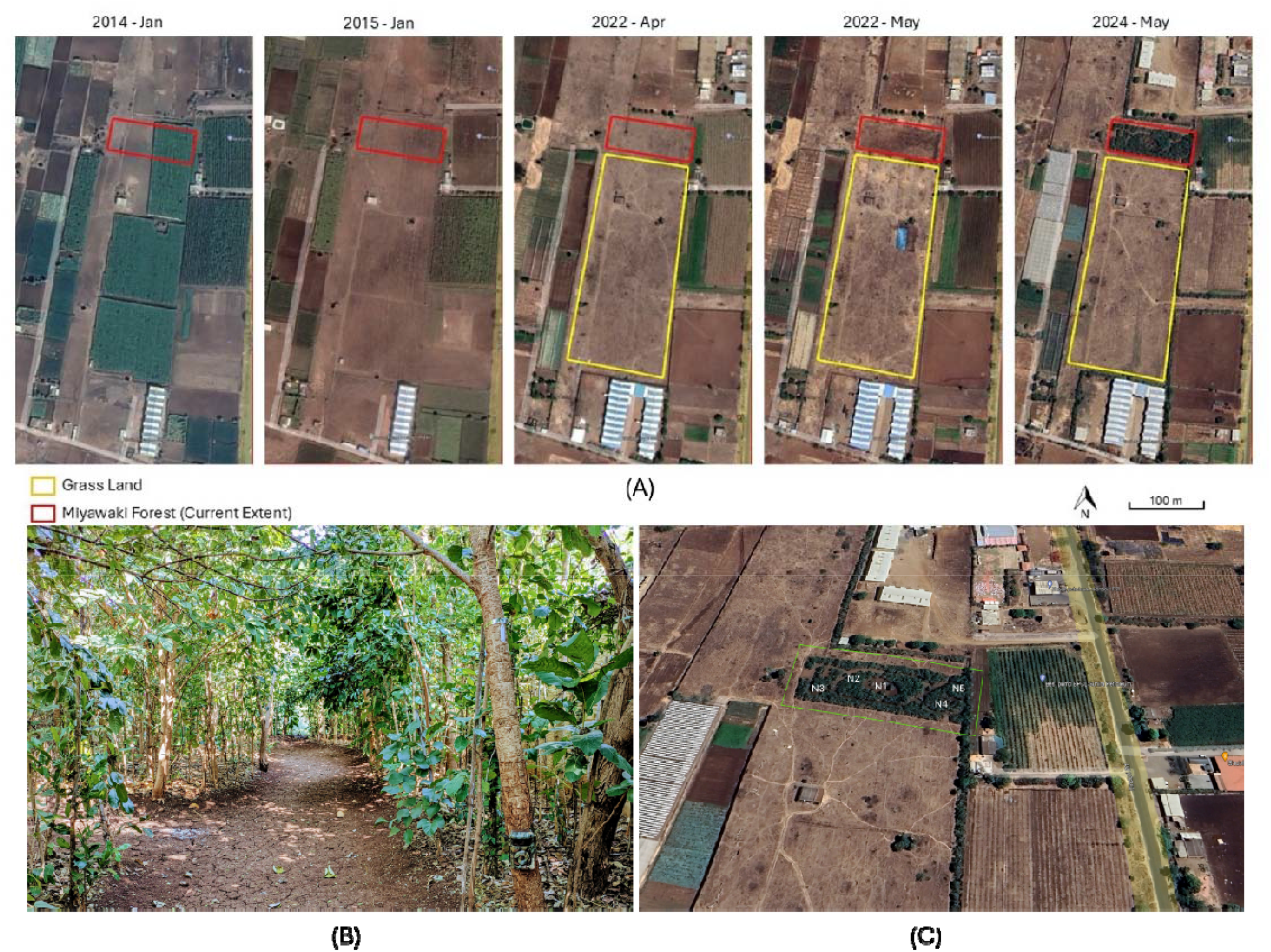
Visual representation of the Miyawaki forest site (A) Google Earth time series illustrating land-cover trajectories at the Miyawaki restoration and adjacent unmanaged grassland in Mohadi, Nashik (2014-2024). Satellite image showing location of Miyawaki forest within the broader landscape (B) Ground-level photograph of the two-year-old Myawaki forest. (C) Satellite image (Google Earth) showing the five grids designated for soil sampling N1, N2, N3, N4 and N5.

### Vegetation structure and soil nutrient dynamics of Miyawaki forest

Tree species selection for Miyawaki plantation followed the original Miyawaki framework. A combination of intensive field surveys of old growth forests in the neighbourhood to document native species composition and their relative abundance and of historical and contemporary Forest Department inventories led to the identification of tree-species for the Miyawaki plantation. The identified tree-species comprised a highly diverse assemblage of native woody species (99 species with a total of 16,626 saplings), grouped into five categories: one core combination of taxa most frequently observed in old growth forests and four additional groups corresponding to vertical forest strata of a natural tropical forest with emergents comprising (11.7%), shrubs (9%), sub-trees (39.9%) and trees (39.4%). The forest’s plant density was maintained at 3.33/m^2^ (supplementary table 1). The planted species consortia formed a multi-stratal architecture, integrating canopy-forming trees (*Ficus* spp. *Mangifera indica, Tectona grandis, Terminalia* spp.), nitrogen fixing legumes (*Albizia, Millettia, Senegalia*), fruiting and nectariferous taxa (*Syzygium, Madhuca, Ficus* spp.), along with numerous timber-yielding and medicinally important species (*Dalbergia, Boswellia, Aegle, Terminalia*). This stratified, frequency informed selection was designed to replicate natural forest architecture, enhance ecological realism and promote structural complexity during early forest development.

Soil sampling was conducted two years after forest establishment during wet season, followed by repeat sampling six months later in the dry season. The soil nutrient analysis reveals significant differences in nutrient dynamics between grassland (C1, C2 and C3) and Miyawaki forest setup (N1, N2, N3, N4 and N5) particularly in the dry season (Table 1, Figure 1(C)). For instance, organic carbon content, a key indicator of soil fertility, remained similar in wet season across land-use types (around 0.6%) but in dry season it increased dramatically in forest with 2.7% - 5.07% compared to grassland with 1.98% to 2.25%. An increase in soil carbon in dry season could be attributed to the increased litterfall among deciduous Indian trees in these conditions. Contrary to this, in dry season, grassland had higher nitrogen concentrations (194.43 to 213.25 kg/ha) as compared to the forest (131.71 to 188.16 kg/ha) likely reflecting reduced vegetation growth and microbial immobilization in grassland as compared to more active nitrogen assimilation and plant uptake in forest soils.

**Table 1.** Nutrient analysis of soil samples.

| Sl no. | Parameters | Units | Wet season |  |  |  |  |  |  |  | Dry season |  |  |  |  |  |  |  |
| --- | --- | --- | --- | --- | --- | --- | --- | --- | --- | --- | --- | --- | --- | --- | --- | --- | --- | --- |
|  |  |  | C1 | C2 | C3 | N1 | N2 | N3 | N4 | N5 | C1 | C2 | C3 | N1 | N2 | N3 | N4 | N5 |
| 1 | pH |  | 7.73 | 6.77 | 7.05 | 7.47 | 7.6 | 7.61 | 7.67 | 7.77 | 6.82 | 7.64 | 6.93 | 7.285 | 7.58 | 7.67 | 7.74 | 7.92 |
| 2 | Electrical Conductivity | dS/m | 0.14 | 0.06 | 0.13 | 0.1 | 0.14 | 0.13 | 0.16 | 0.21 | 0.28 | 0.5 | 0.34 | 0.4 | 0.48 | 0.49 | 0.49 | 0.44 |
| 3 | Organic Carbon | % | 0.66 | 0.6 | 0.62 | 0.63 | 0.66 | 0.63 | 0.57 | 0.63 | 1.98 | 2.25 | 2.11 | 2.7 | 3.66 | 5.07 | 2.7 | 2.37 |
| 4 | Nitrogen | kg/ha | 119.17 | 94.08 | 100.62 | 106.62 | 125.44 | 119.17 | 94.08 | 106.62 | 194.43 | 213.25 | 213.14 | 163.07 | 188.16 | 131.71 | 156.8 | 144.26 |
| 5 | Phosphorus |  | 86.01 | 94.4 | 92.2 | 90.71 | 87.44 | 84.59 | 80.33 | 87.01 | 90.99 | 91.28 | 91.1 | 83.03 | 100.52 | 82.03 | 94.83 | 81.61 |
| 6 | Potash |  | 35.48 | 20.97 | 29.22 | 66.39 | 77.41 | 67.33 | 64.78 | 73.38 | 600.77 | 830.59 | 815.68 | 1143.74 | 998.59 | 1135.68 | 1220.35 | 883.01 |
| 7 | Calcium | Meq/100g | 21.2 | 13.5 | 19.35 | 17 | 16.7 | 19.2 | 26.3 | 23.4 | 20.1 | 27.3 | 26.7 | 21 | 21.9 | 31.8 | 40.8 | 32.8 |
| 8 | Magnesium |  | 10 | 11.5 | 11.3 | 13.2 | 13.9 | 13.7 | 20 | 15.6 | 18.3 | 17.4 | 17.23 | 28.2 | 20.6 | 15 | 13.4 | 18.2 |
| 9 | Sulphur | PPM | 6.09 | 5.67 | 6.81 | 6.17 | 6.33 | 6.33 | 7.15 | 6.99 | 10.36 | 12.09 | 11.08 | 11.68 | 12.58 | 11.02 | 12.25 | 10.94 |
| 10 | Zinc |  | 1.47 | 1.08 | 1.22 | 2.26 | 2.28 | 1.91 | 1.34 | 1.39 | 0.66 | 2.55 | 1.78 | 2.47 | 1.17 | 0.42 | 0.78 | 0.55 |
| 11 | Manganese |  | 2.55 | 6.18 | 3.34 | 5.53 | 3.29 | 4.47 | 3.01 | 3.42 | 14.57 | 62.21 | 20.56 | 18.96 | 26.28 | 27.04 | 81.28 | 15.79 |
| 12 | Iron |  | 5.98 | 14.92 | 6.45 | 4.98 | 2.94 | 4.42 | 2.2 | 2.19 | 20.97 | 15.99 | 19.58 | 5.83 | 5.73 | 3.97 | 2.14 | 4.85 |
| 13 | Copper |  | 3.15 | 2.41 | 2.5 | 2.3 | 1.35 | 1.86 | 2.72 | 2.92 | 4.05 | 5.14 | 4.78 | 3.17 | 0.46 | 4.2 | 4.95 | 2.53 |

Available potash in wet season forest plots ranged between 64.78 to 77.41 mg/kg and grassland had only 20.97 to 35.48 mg/kg. Strikingly, in dry season potash had substantially increased in forest soils (883.01 to 1220.35 mg/kg) far surpassing grasslands (600.77 to 830.59 mg/kg) most likely due to leaf litter and rhizodeposition. Other macronutrients like calcium, magnesium and sulphur were markedly higher in dry season forest soils as compared to wet season, indicating active mineralization in dry season forest. Micronutrients like manganese and copper were higher in dry season forest soils, while zinc and iron showed contrasting seasonal trends across land uses. Soil pH largely remained near neutral across all conditions, while electrical conductivity increased in dry season with forest soil showing values of 0.49 dS/m, suggesting elevated ion concentration due to increased evapotranspiration.

### Sequencing depth and rarefaction

The rarefaction curve demonstrated that all samples reached near asymptotic plateaus, indicating that sequencing depth was sufficient to capture the majority of bacterial diversity present (Figure 2). Total sequencing generated 17.7 million paired-end reads, with an average of 0.49 million paired reads per sample (range: 0.22 – 0.97 million paired reads), with Q30 scores ranging from 90.61% to 94.85%. Forest and grassland soils exhibited comparable sequencing coverage, suggesting that the observed differences in diversity metrics primarily reflect ecological variations rather than technical bias (Table 2).

**Table 2.** Metadata of soil samples used in the present study.

| Sample ID | Ecology | Season | R1 Reads | R1 Bases | R1 Q30 (%) | R2 Reads | R2 Bases | R2 Q30 (%) | Paired Reads | Total Bases | Avg. Read Length | Mean Q30 (%) | Mean GC (%) | Accession ID |
| --- | --- | --- | --- | --- | --- | --- | --- | --- | --- | --- | --- | --- | --- | --- |
| N1aW | Forest | Wet | 932416 | 280657216 | 94.92 | 932416 | 280657216 | 88.6 | 932416 | 561314432 | 301 | 91.76 | 57.89 | SRR35758316 |
| N1bW | Forest | Wet | 433230 | 130402230 | 96.8 | 433230 | 130402230 | 92.5 | 433230 | 260804460 | 301 | 94.65 | 57.63 | SRR35758285 |
| N1cW | Forest | Wet | 276970 | 83367970 | 96.66 | 276970 | 83367970 | 91.97 | 276970 | 166735940 | 301 | 94.315 | 57.58 | SRR35758284 |
| N2aW | Forest | Wet | 923841 | 278076141 | 95.16 | 923841 | 278076141 | 89.23 | 923841 | 556152282 | 301 | 92.195 | 57.17 | SRR35758315 |
| N2bW | Forest | Wet | 522749 | 157347449 | 96.77 | 522749 | 157347449 | 92.49 | 522749 | 314694898 | 301 | 94.63 | 57.365 | SRR35758283 |
| N2cW | Forest | Wet | 612076 | 184234876 | 96.76 | 612076 | 184234876 | 92.72 | 612076 | 368469752 | 301 | 94.74 | 56.995 | SRR35758282 |
| N3aW | Forest | Wet | 844651 | 254239951 | 94.99 | 844651 | 254239951 | 89.28 | 844651 | 508479902 | 301 | 92.135 | 57.48 | SRR35758304 |
| N3bW | Forest | Wet | 520632 | 156710232 | 96.87 | 520632 | 156710232 | 92.84 | 520632 | 313420464 | 301 | 94.855 | 57.375 | SRR35758281 |
| N3cW | Forest | Wet | 588290 | 177075290 | 96.76 | 588290 | 177075290 | 92.71 | 588290 | 354150580 | 301 | 94.735 | 57.44 | SRR35758314 |
| N4aW | Forest | Wet | 978986 | 294674786 | 94.93 | 978986 | 294674786 | 86.29 | 978986 | 589349572 | 301 | 90.61 | 57.885 | SRR35758293 |
| N4bW | Forest | Wet | 678433 | 204208333 | 96.85 | 678433 | 204208333 | 92.51 | 678433 | 408416666 | 301 | 94.68 | 57.8 | SRR35758313 |
| N4cW | Forest | Wet | 591203 | 177952103 | 96.77 | 591203 | 177952103 | 92.54 | 591203 | 355904206 | 301 | 94.655 | 57.795 | SRR35758312 |
| N5aW | Forest | Wet | 893753 | 269019653 | 95.19 | 893753 | 269019653 | 88.7 | 893753 | 538039306 | 301 | 91.945 | 57.975 | SRR35758286 |
| N5bW | Forest | Wet | 624732 | 188044332 | 96.77 | 624732 | 188044332 | 91.53 | 624732 | 376088664 | 301 | 94.15 | 57.715 | SRR35758311 |
| N5cW | Forest | Wet | 597810 | 179940810 | 96.75 | 597810 | 179940810 | 92.45 | 597810 | 359881620 | 301 | 94.6 | 58.075 | SRR35758310 |
| NC1W | Grassland | Wet | 832223 | 250499123 | 95.27 | 832223 | 250499123 | 89.07 | 832223 | 500998246 | 301 | 92.17 | 58.3 | SRR35758309 |
| NC2W | Grassland | Wet | 885824 | 266633024 | 95.34 | 885824 | 266633024 | 88.73 | 885824 | 533266048 | 301 | 92.035 | 58.89 | SRR35758308 |
| NC3W | Grassland | Wet | 562854 | 169419054 | 96.8 | 562854 | 169419054 | 92.19 | 562854 | 338838108 | 301 | 94.495 | 58.37 | SRR35758307 |
| N1aD | Forest | Dry | 220599 | 66400299 | 93.78 | 220599 | 66400299 | 90.58 | 220599 | 132800598 | 301 | 92.18 | 57.5 | SRR35758306 |
| N1bD | Forest | Dry | 225264 | 67804464 | 93.62 | 225264 | 67804464 | 90.29 | 225264 | 135608928 | 301 | 91.955 | 57.52 | SRR35758305 |
| N1cD | Forest | Dry | 282010 | 84885010 | 94.68 | 282010 | 84885010 | 91.2 | 282010 | 169770020 | 301 | 92.94 | 57.465 | SRR35758303 |
| N2aD | Forest | Dry | 275125 | 82812625 | 94.54 | 275125 | 82812625 | 91.49 | 275125 | 165625250 | 301 | 93.015 | 56.74 | SRR35758302 |
| N2bD | Forest | Dry | 325195 | 97883695 | 94.65 | 325195 | 97883695 | 91.07 | 325195 | 195767390 | 301 | 92.86 | 57.05 | SRR35758301 |
| N2cD | Forest | Dry | 666349 | 200571049 | 94.39 | 666349 | 200571049 | 90.91 | 666349 | 401142098 | 301 | 92.65 | 57.135 | SRR35758300 |
| N3aD | Forest | Dry | 293948 | 88478348 | 94.43 | 293948 | 88478348 | 91.29 | 293948 | 176956696 | 301 | 92.86 | 57.1 | SRR35758299 |
| N3bD | Forest | Dry | 271364 | 81680564 | 94.64 | 271364 | 81680564 | 91.2 | 271364 | 163361128 | 301 | 92.92 | 57.175 | SRR35758298 |
| N3cD | Forest | Dry | 243722 | 73360322 | 94.11 | 243722 | 73360322 | 90.97 | 243722 | 146720644 | 301 | 92.54 | 57.25 | SRR35758297 |
| N4aD | Forest | Dry | 271407 | 81693507 | 94.46 | 271407 | 81693507 | 90.51 | 271407 | 163387014 | 301 | 92.485 | 57.77 | SRR35758296 |
| N4bD | Forest | Dry | 249519 | 75105219 | 94.35 | 249519 | 75105219 | 90.66 | 249519 | 150210438 | 301 | 92.505 | 58.07 | SRR35758295 |
| N4cD | Forest | Dry | 305410 | 91928410 | 94.16 | 305410 | 91928410 | 90.69 | 305410 | 183856820 | 301 | 92.425 | 57.995 | SRR35758294 |
| N5aD | Forest | Dry | 343841 | 103496141 | 94.66 | 343841 | 103496141 | 90.74 | 343841 | 206992282 | 301 | 92.7 | 57.75 | SRR35758292 |
| N5bD | Forest | Dry | 305580 | 91979580 | 94.03 | 305580 | 91979580 | 90.49 | 305580 | 183959160 | 301 | 92.26 | 57.565 | SRR35758291 |
| N5cD | Forest | Dry | 288467 | 86828567 | 94.21 | 288467 | 86828567 | 90.76 | 288467 | 173657134 | 301 | 92.485 | 57.91 | SRR35758290 |
| NC1D | Grassland | Dry | 294654 | 88690854 | 94.49 | 294654 | 88690854 | 90.11 | 294654 | 177381708 | 301 | 92.3 | 58.35 | SRR35758289 |
| NC2D | Grassland | Dry | 281410 | 84704410 | 94.22 | 281410 | 84704410 | 90.16 | 281410 | 169408820 | 301 | 92.19 | 58.28 | SRR35758288 |
| NC3D | Grassland | Dry | 316552 | 95282152 | 94.42 | 316552 | 95282152 | 90.48 | 316552 | 190564304 | 301 | 92.45 | 58.21 | SRR35758287 |

**Figure 2.**
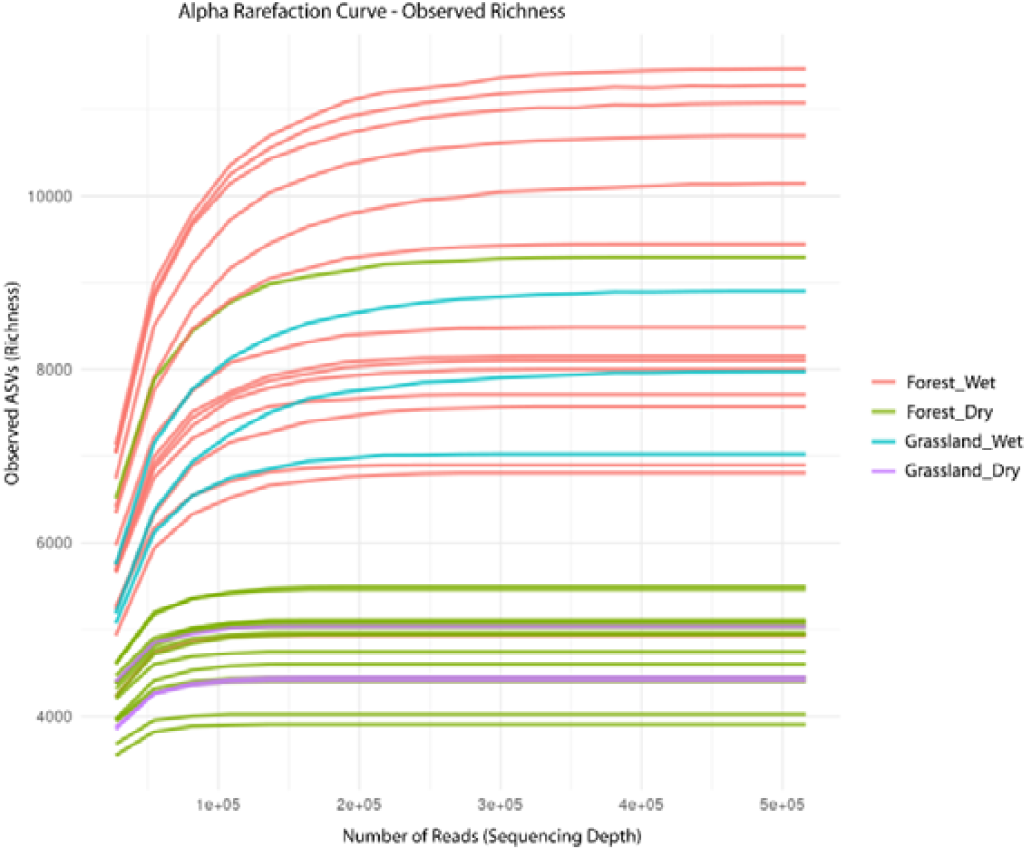
Rarefaction curves illustrating soil bacterial diversity across ecosystems and seasons. The curves represent number of amplicon sequence variants (ASVs) as a function of sequencing depth estimating bacterial richness.

### Alpha diversity patterns of soil microbial communities across habitats and seasons

Bacterial richness, as depicted by Observed and Chao1 indices was significantly higher (p = 0.00015) in forest soils, particularly in wet conditions as compared to the grasslands (Figure 3). Greater plant diversity and stable organic matter inputs in forest suggests that forest provides favourable niche for a diversity of bacterial taxa than grassland. Seasonal richness decline was observed in dry season forest soils, possibly indicating transient stress despite artificial watering. Additionally, Shannon (*p* = 0.0057) diversity index significantly highlighted that bacterial communities also vary in evenness and dominance structure across seasons in a particular habitat. In contrast, grassland had fewer dominant taxa in the community as implicated by lower richness and evenness. These findings provide a useful ecological baseline to the forest comprising of richer, diverse and seasonally dynamic bacterial communities.

**Figure 3.**
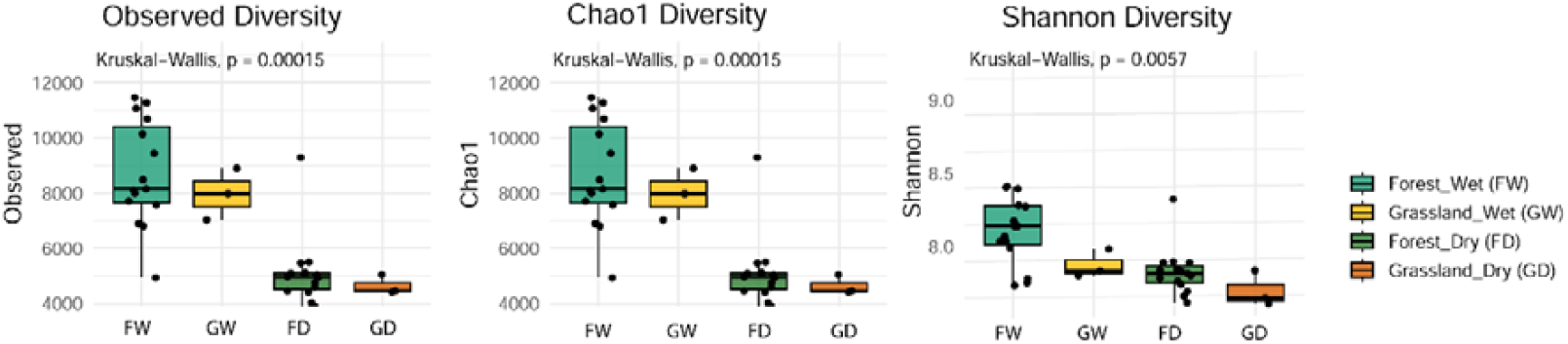
Comparative analysis of soil bacterial diversity (A) Observed ASVs, (B) Chao1 diversity and (C) Shannon Diversity index across forest and grassland in wet and dry seasons.

### Beta diversity of soil microbial communities across habitats and seasons

Principle Coordinate Analysis (PCoA) based on Bray-Curtis dissimilarity revealed clear structuring of taxonomic diversity by habitat, however, seasonal differentiation was not evident within the habitat (Figure 4). The first two PCoA axes explained 23.7% and 16.4% of the total variation and PERMANOVA analysis (adonis2) revealed a significant separation on bacterial community composition (F = 5.57, R^2^ = 0.34, p = 0.001). However, functional beta diversity based on predicted bacterial metabolic functions showed a cluster for forest samples, wherein, the dry season grassland samples were also partially overlapping indicating functional convergence. PCoA explained 51.1% and 19.6% variation and PERMANOVA confirmed significance of the analysis (F = 5.43, R^2^ = 0.34, p = 0.002).

**Figure 4.**
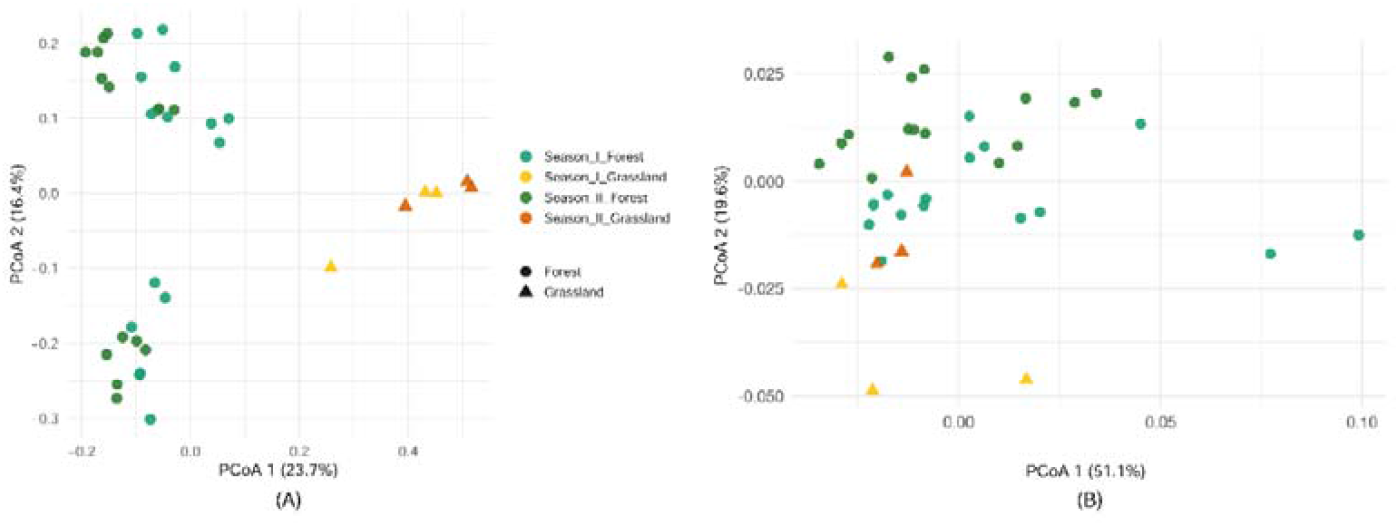
Beta diversity of soil bacterial communities across ecosystems and seasons. (A) Taxonomic beta diversity based on community composition and (B) functional beta diversity inferred from predicted metabolic functions of bacterial communities.

### Bacterial functional shifts across ecologies and seasons

Functional prediction of bacterial communities performed using PICRUSt2 inferred 7,688 Kyoto Encyclopedia of Genes and Genomes (KEGG) Orthologs (KOs) from 16S rRNA metagenome data. Differential abundance analysis was performed using DESeq2 and the predicted functional profiles across all samples were visualized using volcano plots to highlight differential KOs across key pairwise comparisons (Figure 5). These plots depict the statistical significance (–log10 adjusted p-value) against the magnitude of change (log2FoldChange) for each KO illustrating patterns of enrichment or depletion of KOs across different comparisons. Notably, seasonal changes within forest depicts fewer KOs showing significant shifts, indicating a relatively stable functional profile across seasons in the forest environment. In contrast, significantly enriched KOs in grassland dry season suggest functional stress responses under dry, non-watered conditions. Furthermore, forest versus grassland comparisons showed more dispersed volcano patterns, reflecting larger functional divergences between land use types.

**Figure 5.**
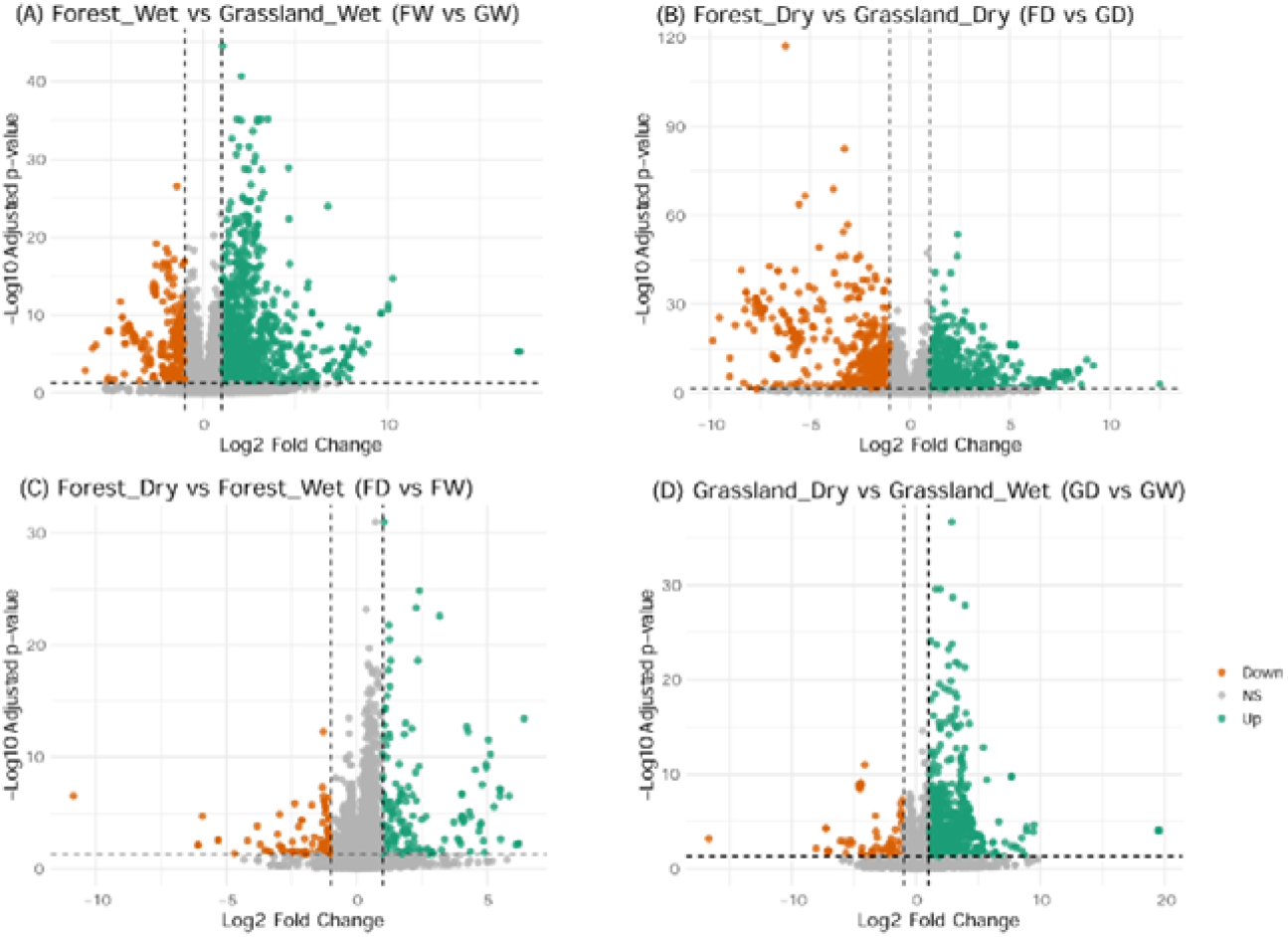
Volcano plots representing differential abundance of KEGG Orthology (KO) functions across conditions analysed using DESeq2. Down: downregulated, NS: non-significant and Up: upregulated KOs.

Out of 7,688 KOs, 1,427 were meeting the statistical significance and abundance thresholds (mentioned in methods). These significantly varying KOs were distributed across 82 KEGG pathways (Figure 6, supplementary table 2). Majority of which mapped to core metabolic pathways (map01100, n=478 KOs) indicating widespread metabolic reprogramming across environmental gradients. Biosynthesis of secondary metabolites (map01110, n=172 KOs) and microbial metabolism in diverse environments (map01120, n=163 KOs) reflecting adaptive microbial responses to changing environments were noted. Further, two component systems (map02020, n=72 KOs), ABC transporters (map02010, n=69) reflecting altered microbial sensing and transport regulation were observed. In addition to these categories, diverse set of environment and energy linked pathways like photosynthesis (map00195, n=47 KOs), methane metabolism (map00680, n=42 KOs), aromatic compound degradation (map01220, n=34 KOs) were recorded. Further, the observed quorum sensing (map02024, n=25 KOs) and biofilm formation pathways (map02026, map05111, map02025, n=26 KOs) points towards changes in microbial communication. Together these patterns are indicative of functional reorganization of microbial communities beyond metabolism, encompassing environmental sensing, signalling and adaptation.

**Figure 6.**
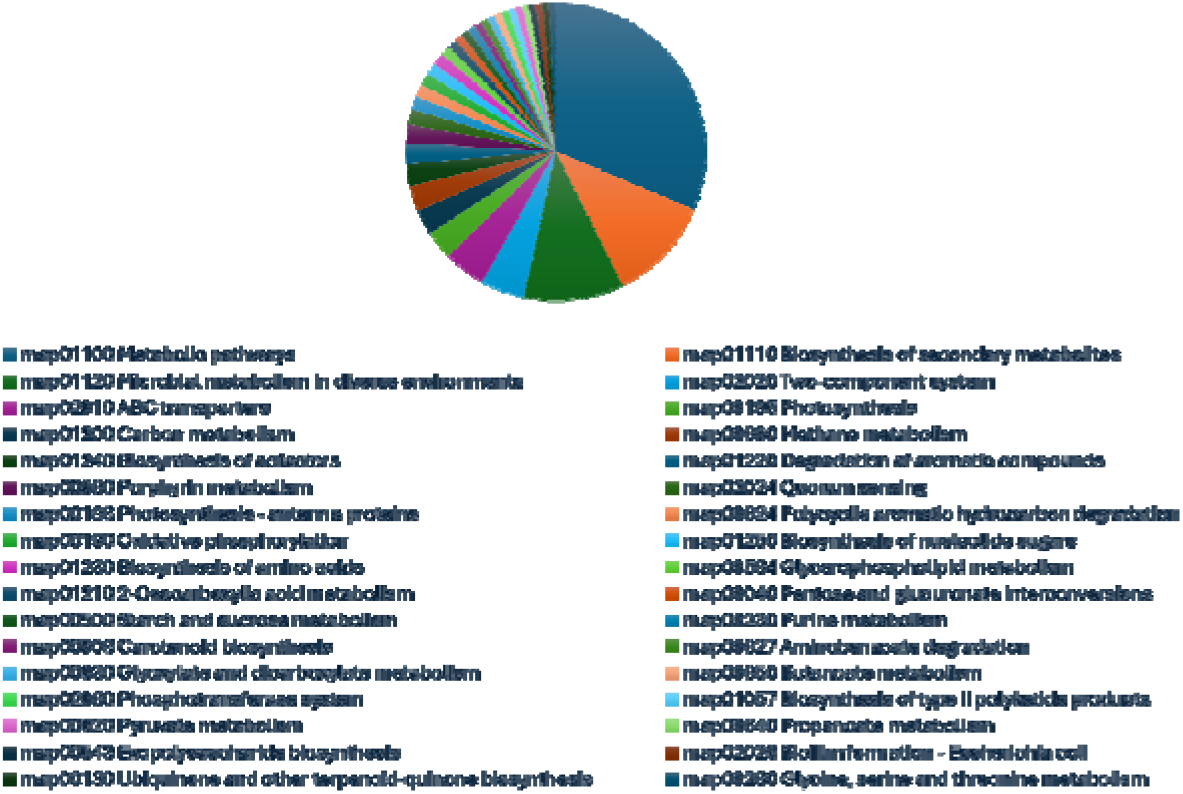
Pie-chart represents distribution of pathways map IDs associated with KO IDs (for clarity representing only map IDs having > 10 KO IDs) showing significant upregulation or downregulation across ecosystem types in dry season.

### Differential regulation of gene clusters reflects habitat specific functional remodelling

Shifts with the most substantial divergence were in Miyawaki forest versus grassland in the dry season, comprising 66 upregulated and 43 downregulated KOs (supplementary table 2). Forest in dry season showed strong enrichment for anaerobic and redox-flexible respiratory pathways including Na^+^-transporting NADH: ubiquinone oxidoreductase complex (*nqrABCDEF* (K00346-K00351)) ensuring enhanced energy generation and pH homeostasis in micro-anoxic niches of decomposing litter. Nitrate/nitrite reduction (*narP, fdnHI* (K07685, K08349-K08350)), arsenite oxidation system (*aoxAB* (K08355-K08356)), selenate/chlorate reduction complex (*serAB, clrAB* (K17050-K17051)) expanding respiratory flexibility in fluctuating redox microsites and formate hydrogenlyase complex (*hycBCDEFGH* (K15827-K15834)) suggests hydrogen-based syntrophy during anaerobic litter decomposition. Catabolism of complex and recalcitrant carbon was elevated, including aromatic and hydrocarbon degradation modules (*4CL* (K01904), *nidAB* (K11943-K11944)) toluene monooxygenase complex (*tmoABDE* (K15760-K15764)) and sulfur-rich polymer breakdown systems (*ARSA/B, IDS* (K01134-K01136)) involved in hydrolysis of plant cell wall polysaccharides.

Additionally, in Miyawaki forest ecosystem stress adaptation was marked by sporulation (*cotVWXYZ* [K06340–K06343], *rapABDEGHIK* [K06359– K06360, K06362-K06363, K06365-K06367, K06369], universal stress proteins ensuring population survival through seasonal fluctuations. EPS/biofilm production genes (*algE* [K16081], *algK* [K19292], *algX* [K19293], and *algF* [K19296], *exoM* [K16556], *exoA* [K16557], *exoI* [K16561], *exoX* [K16565], *exoQ* [K16567], *epsK* [K19418], *epsA* [K19420], *epsC* [K19421], *epsE* [K19423], *epsF [*K19424], *epsI* [K19426], *epsN* [K19430] promote colonization of plant debris. Nutrient-scavenging systems, including siderophores (*iucABC* [K03894–K03896]), metal transporters (*mtsABC* [K11704-K11706], *mntCB* [K11601-K11602], *sitABCD* [K11604-K11607]), and amino acid transporters (*artJI* [K09996-K09997], *attA1A2BC* [K11077-K11081]), were also upregulated in Miyawaki for survival in competitive litter microhabitats.

In contrast, grassland in dry season maintained high abundance of oxygenic phototrophic genes, while forest dry showing downregulation of photosystem I/II (*psaABCDEFIJKLMX* [K02689–K02702], *psbABCDEFHIJKLMNOPTUVXYZ* [K02703–K02724]), cytochrome b6-f / ferredoxin-plastocyanin system (*petABCDEFGHLM* [K02634-K02643]) phycobilisomes (*apcABCDEF* [K02092–K02097], *cpcABCDEFG* [K02284–K02290], *cpeABCRSTUYZ* [K05376–K05378, K05381-K05386]) and carbon concentrating mechanisms (*ccmKLMNO* [K08696–K08700]). These changes reflect a shift from light driven primary production towards heterotrophic metabolism under canopy shading. Grassland soils also display upregulation of carotenoid biosynthesis genes (*crtO* [K02292], *crtP* [K02293], *crtR* [K02294], *crtISO* [K09835], *crtW* [K09836], *crtC* [K09844], *crtD* [K09845], *crtF* [K09846]) consistent with adaptation to oxidative stress. Light harvesting suppression and litter associated colonization functions persisted in forest wet season.

### Enrichment of Plant Growth Promoting Rhizobacteria (PGPRs) across habitat

PGPRs facilitate plant growth through multiple mechanisms, including nitrogen fixation, phosphate solubilization, phytohormone modulation and siderophore-mediated micronutrient acquisition (Compant et al., 2010; Kloepper et al., 2004). We examined the distribution of PGPRs in forest and grassland soils. A diverse array of PGPR genera like: *Bacillus, Bradyrhizobium, Flavobacterium, Mesorhizobium, Microbacterium, Paenibacillus, Pseudomonas* and *Variovorax* were enriched in forest with p-value < 0.05 (Figure 7). The significant enrichment of well-characterized PGPRs in the Miyawaki forest soil indicates that dense native vegetation, altered environment, apart from just soil management strategies for forest establishment together influencing the selection of the microbiome that can facilitate an early forest establishment. These taxa are known to enhance nutrient acquisition through biological nitrogen fixation, phosphorus solubilization and synthesis of phytohormones that promote root growth and alleviate abiotic stresses (Backer et al., 2018; Glick, 2012; Olanrewaju et al., 2017). Together, these functional attributes imply that the microbial shifts in Miyawaki forest constitute foundational support for the establishment of improved nutrient cycling and progressive development of a stable forest soil ecosystem.

**Figure 7.**
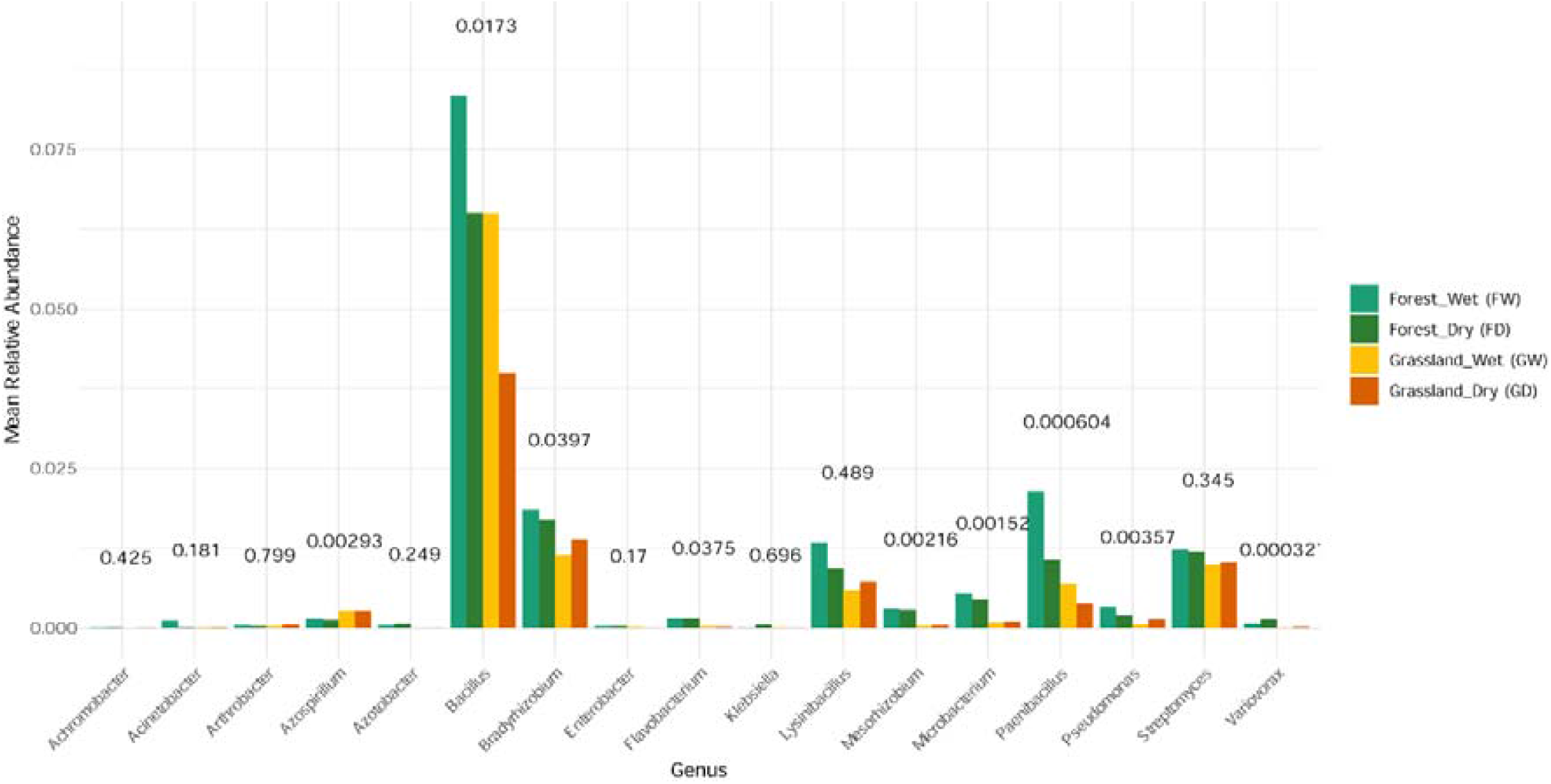
Relative abundances of selected plant growth-promoting rhizobacteria (PGPR) groups in forest and grassland soils with corresponding p-values on the top of each bar.

## Discussion

Recent evaluations of the benefits of Miyawaki method for ecosystem restoration have highlighted limited mechanistic understanding due to lack of empirical data with replicated study designs, appropriate controls, comparisons with other restoration methods and temporally resolved data (Morales et al., 2025). In the present study, we implemented replicated soil sampling strategy across multiple Miyawaki forest sites in addition to the neighbouring grassland as a reference system and examined seasonal dynamics by sampling during wet and dry seasons. The study aimed to characterise soil bacterial communities in the two ecosystems as soil microbiome acts as a sensitive indicator of ecosystem development in response to the vegetation driven changes and microenvironmental conditions (Bardgett & Van Der Putten, 2014).

### Vegetation driven structuring of microbial communities

The Miyawaki forest investigated in the present study represents an emerging functionally heterogeneous assemblage. The 99 species integrate multiple functional guilds like: evergreen and deciduous taxa, nitrogen fixing legumes, deep and shallow rooted species, as well as fruit bearing taxa. Such complementary traits accelerate biomass accumulation, buffer environmental variability and increase probability of establishing self-sustaining nutrient and hydrological cycles (Díaz et al., 2007; Poorter et al., 2021). The integration of fast-growing pioneers (*Albizia procera* and *Senegalia ferruginea*) contribute to early-stage nutrient cyclers, canopy formation and rapid biomass turnover. The coexistence of evergreen species (*Syzygium cumini, Ficus benghalensis, Mangifera indica*) ensures year-round canopy cover, microclimatic buffering and continuous litter inputs, whereas, deciduous species (*Albizia lebbeck, Bauhinia variegata, Boswellia serrata*) generate temporal variability in resource availability. Further, the litter deposition and soil organic matter enhancement, macro-pore development due to soil fauna, diverse range of rooting depths may improve water infiltration and resilience to seasonal droughts, but may also deplete soil moisture resulting in shallow ground water due to the infiltration-evapotranspiration trade-off (Bruijnzeel et al., 2025; Ilstedt et al., 2007; Krishnaswamy et al., 2018).

Furthermore, multiple *Ficus* spp., *Syzygium* spp. and other fruiting trees provide fruits and nectar resources that attract birds, bats and pollinator guilds thereby, enhancing trophic interactions establishing seed dispersal mechanisms (Shanahan, 2000). Hence, such a species rich approach to achieve functional complementarity can drive rapid enhancement of multifunctional ecosystem services in urban and peri-urban context. Further, the remotely sensed time-series of the warehouse site also highlights the capacity of the Miyawaki method to rapidly alter the above-ground vegetation structure within a short temporal window, transitioning to a dense, multi-layered canopy in a span of just two years.

Soil microbial diversity and function are largely influenced by vegetation cover as it governs habitat structure, resource inputs and microclimatic buffering (Baldrian, 2017; Delgado-Baquerizo et al., 2018; Fierer & Jackson, 2006). The dense canopy in a forest provides nutrient rich complex litter and shaded environment supporting higher microbial richness and evenness particularly in wet season. In contrast, open system grasslands is characterized by high sun irradiance imposing physiological stress on soil microbes, often altering microbial activity and community structure in response to moisture limitation (Evans & Wallenstein, 2012). The present forest vs grassland soils depicted functional tailoring of the microbial assemblages across the habitats and seasons. Dry season depicted the strongest divergence with specialized microbial communities in both the habitats. Seasonal drying amplifies the gradients of oxygen, nutrient, osmotic, thermal and redox stresses leading to microbial competition and favouring the stress-resilient community with nutrient uptake, sporulation, biofilm formation abilities (Baldrian, 2017; Keiluweit et al., 2018). On the other hand, in the monsoon season the continuous moisture availability dilutes the heterogenous stress gradients from the dry season, buffering microbial activity (Bouskill et al., 2013; Placella et al., 2012). Collectively, these ecological contrasts derived by vegetation and seasonal impact magnify community level adaptations ultimately featuring functional contrasts amongst Miyawaki forest and grassland soils.

### Redox flexibility and micro-anoxia in forest soil

In forest soils during the dry season, dense canopy and continuous litter accumulation creates a less ventilated and cooler forest floor than the adjacent open grassland. This leads to formation of microsites with patchy oxygen availability creating fine-scale heterogeneity, influencing soil microbial activity and community structure (Silver et al., 1999). Microbial communities exhibited a shift towards redox flexibility and anaerobic metabolism, supported by enrichment of the Na□-pumping respiratory entry module *nqrABCDEF*, nitrate/nitrite regulators (*narP*), formate dehydrogenase (*fdnHI*), formate hydrogenlyase (*hycBCDEFGH*) and alternate electron accepting systems including arsenite oxidation (*aoxAB*) and selenate/chlorate reduction (*serAB, clrAB*). These adaptations allow energy conservation under oxygen-limited conditions and underscores how forest soils harbor micro-anoxic niches despite being overall aerated. Indeed, forest soils are recognized as mosaics of oxic and anoxic microsites created by aggregate structures and higher litter inputs (Keiluweit et al., 2018). Hence, micro-scale anoxia drives nitrate reduction, hydrogen turnover and alternative electron acceptor use, extending metabolic breadth of forest microbiomes.

### Forest microbiomes as detritus specialists

Forest litter is rich in lignin and phenolics and microbial communities in forest soils are specialized in their degradation (Wilhelm et al., 2019). A hallmark of the forest microbiome was enrichment of aromatic degrading and polymer cleaving enzymes such as toluene monooxygenase (*tmoABDE*), oxygenases (*ligXa*), PAH dioxygenases (*nidAB*), 4-coumarate: CoA ligase (*4CL*), sulfatases (*ARSAB, IDS*). These enzymes initiate oxidation and funnelling of lignin-derived aromatics and hydrolyse sulphated polysaccharides from the tree sheds. Sulfatase activity has been linked to late-stage decomposition of chemically complex litter, contributing to complex carbon turnover (Kunito et al., 2022; Ponder & Eivazi, 2008). Collectively, forest microbial community genes are enriched for deep litter degradation functions.

### Microbial community dynamics and resource competition in forest soils

Forest soils also displayed signatures of microbial persistence strategies by enrichment of exopolysaccharide (EPS) and biofilm gene clusters (*alg, exo* and *eps* operons). EPS enhances water retention in micropores and alters pore-scale water distribution creating highly hydrated microenvironment and its shrink swell behaviour buffers soil against rapid drying (Guo, 2018; Or et al., 2007; Roberson & Firestone, 1992). Further, EPS cements the soil particles resulting in aggregate stability ensuring soil remains porous rather than compacted while biofilm ensures interspecies interactions (Chenu, 1995; Flemming & Wuertz, 2019). Additionally, biofilms are increasingly recognized as a mediator of healthy plant-microbiome interactions, nutrient exchange and protection against pathogens (Compant et al., 2010; Rudrappa et al., 2008).

Although, the forest floors are enriched in organic matter, yet, much of the nutrient pool is chemically immobilized with recalcitrant litter polymers complexed with humic substances (Cleveland et al., 2003; Kuzyakov & Blagodatskaya, 2015). Hence, microbial competition for limiting resources is also a feature of the forest soils. For instance, the bioavailability of iron, an essential cofactor for respiration and oxidative enzymes, is poor, driving selection for microbial nutrient scavenging strategies. Thus, to overcome nutrient scarcity, siderophore biosynthesis clusters (*iucABC*) and high affinity metal transporters (*mtsABC, sitABCD*) were upregulated, correlating with microbial adaptation to micronutrient limited microsites within litter rich soils. Stress adaptation modules further reinforced via sporulation genes (*cot, rap*) and universal stress proteins were abundant, reflecting bet-hedging strategies that enhance survival during seasonal fluctuations (Nicholson et al., 2000). This suggests that the forest microbiome communities organize and evolve themselves to be able to better adapt to seasonal variations making them more robust.

In addition to the stress resilient microbial community, forest soil was also inhabited by diverse PGPRs. These microbes establish mutualistic interactions with plant counterparts which contributes to root system development and enhanced nutrient acquisition. Their prevalence in forest soils depicts that, beyond mere persistence, these bacteria play pivotal role in facilitating growth of host plant and hence, overall vegetation success. PGPRs are known to enhance plant biotic and abiotic stress tolerance, nutrient uptake, thus, supporting ecosystem resilience building (Compant et al., 2010; Kloepper et al., 2004). Extrapolating to the context of Miyawaki forests, once the specialized microflora has developed, PGPRs can sustain nutrient cycling and reduce the need for external inputs or management. Thereby, microbes reinforce ecological stability and economic feasibility of afforestation (Guo, 2018).

### Phototrophic dominance and oxidative stress adaptation in grassland soils

In contrast to forest, grassland microbial communities exhibited strong signatures of phototrophic and oxygenic metabolism. Genes encoding photosystems I and II (*psa* and *psb*), phycobilisomes (*apc, cpc, cpe* operons), cytochrome b6-f complexes, and carbon concentrating mechanisms (*ccm*) were highly abundant. This emphasises the fundamental role of cyanobacteria and other phototrophs in driving primary production under open canopy and high light conditions (Steven et al., 2014). However, in addition to phototrophic gene clusters, upregulation of carotenoid biosynthesis genes in grasslands highlights the prevalence of oxidative stress due to high irradiance and hyperoxidation, supporting the specialization of microbial community in reactive oxygen species (ROS) detoxifying pigments (Latifi et al., 2009).

Collectively, the grassland microbiome has optimized as a light-driven primary producer with oxidative stress combatting, in contrast to the heterotrophic specialization, and persistence strategies that dominate in the forest soils. These contrasting ecological contexts shape microbial communities into distinct taxonomic and functional tailoring. However, it is noteworthy to state that Miyawaki forest was subjected to consistent management including input of organic amendments, watering in dry season etc, unlike the grassland. Yet, diverse ecological filters in forest like: canopy microclimate, complex litter inputs, extensive root systems, compared to open, sun lit, moisture variable grassland were vegetation driven and are not results of direct management interventions. Hence, the functional adaptation of microbial communities to the habitats indicate that these are fundamental ecological strategies and not just management artifacts. Here, the unmanaged grassland represents a microbial baseline characterized by phototrophy and antioxidative defences (Latifi et al., 2009). While, forest represents the other end of the ecological succession, dominated by heterotrophic decomposition, widening up their metabolic spectrum enabling microbes to consume complex litter while withstanding micro-anoxic niches (Baldrian, 2017; Keiluweit et al., 2018). Such shifts are ecologically significant in Miyawaki afforestation models, where microbial trajectories from grassland-like baselines evolve into a forest-like functional reorganization providing a foundation for early-stage soil development inducing forest establishment (Callahan et al., 2016; Guo, 2018).

### Temporal functional reorganization signals early-stage microbial community development in the Miyawaki forests

In the present study, we have observed pronounced functional divergence between wet and dry seasons, which reflects not just the short-term environmental fluctuations due to seasons, but also temporal maturation of soil microbiota concomitant with the forest’s maturation. The dry season sampling conducted nearly six months post-monsoon (wet season) exhibited stronger enrichment for anaerobic respiration, stress response pathways, and aromatic compounds degradation, indicating successional shift towards a metabolically versatile microbial consortium. This pattern is consistent with restoration ecology models where microbial communities evolve towards multifunctionality, higher metabolic complexity as organic matter accumulates and stress amplifies during the forest development (Martin, 2011). The temporal microbial turnover along with the establishment of redox-flexible and stress-adaptive microbial communities parallels with the increasing organic matter accumulation, micro-anoxic niche diversification and canopy closure which are key indicators of ecological maturity (Cai et al., 2025; Sun et al., 2025). Thus, the observed seasonal divergence represents tropical restoration chrono-sequences progressing towar increasingly organized and functionally sturdier biogeochemical processes within a span of six months.

## Conclusion

This study provides a microbial baseline for peri-urban Miyawaki forests, showing how dense tree plantations shape soil bacterial communities differently from grasslands. Seasonal shifts reveal successional reorganization of microbial functional traits, reflecting adaptation to organic-rich, oxygen-limited soils in the forest versus sunlit, hyperoxic conditions in grasslands. Integrating microbial and plant traits offers a framework to assess ecological trajectories of urban afforestation. It is noteworthy that a typical natural forest that contains grasses, shrubs and trees with ample spacing for sunlight streams at some places and dense canopies at others would have the advantage of comprising both grassland and Miyawaki forest bacterio-flora contributing to a highly dynamic and resilient ecosystem. Therefore, our results suggest that a combination of trees, shrubs and grasses instead of a tree-centric approach offers a better alternative to achieve sustainable and resilient urban and peri-urban greening.

## Future directions

We acknowledge that this study captures early snapshot in the development of a Miyawaki forest and does not assess its long-term trajectories. It provides an evidence-based assessment of early belowground trajectories in an engineered urban forest system. However, functional interpretations are derived from taxonomic assignments rather than direct measurements of microbial activity. Hence, increased heterotrophic potential in forest soils should be directly assessed by using carbon turnover and sequestration in forest. Future research incorporating long-term monitoring of integrated, multi-trophic parameters along with the microbial functional capacity, carbon stability and resilience across Miyawaki sites will be critical. Additionally, future greening initiatives may benefit from exploring the usage of soil as inoculum from matured forests to accelerate the functional establishment, enhance stress tolerance and guide early successional trajectories.

## Methods

### Site selection and plantation establishment

The study aimed to investigate soil microbial community composition of a Miyawaki forest established in May 2022, in a 1.5-acre open space at a warehouse site in Mohadi, Nashik, India (20.1352095 °N, 73.8890946 °E). The bacterial community from the Miyawaki forest was compared with that of the nearby reference grassland (20.1349013 °N, 73.8887195 °E). Initially the 10-acre warehouse site was a grape farm which was later replaced with an untended grassland plot for 9 years between 2013-2022. In 2022 when the Miyawaki forest was being developed the designated 1.5-acre space was demarcated into beds of size - 3×50 m each. Pits were dug to a depth of 1.5 m and filled with alternating layers of soil and cattle manure sourced from a local cowshed. Saplings were planted in mixed species groups following Miyawaki method of plantation with an average spacing of 60 cm between individuals to promote dense growth. After planting, each bed was provided with 10-12 cm layer of organic mulch and irrigated. Saplings were watered twice weekly for the first year, after which irrigation was gradually phased out over the next 2.5 years. Post plantation, the only soil treatments involved the addition of mulch to the beds once every few months in the first year.

### Sample collection and nutrient analysis

For the study, we have sampled the forest across two seasons: wet (August 2024) and dry (March 2025). Soil samples were collected from five grids (N1, N2, N3, N4 and N5) (Figure 1) in triplicates across forest and from one random site in triplicates (C1, C2 and C3) from the neighbouring grassland on 30th August 2024. Soil sampling was repeated in triplicates from the same sites in the dry season on 28th March 2025. A total of 36 soil samples were collected in 1.5 kg sterile bags and in 50 mL falcon tubes each. Falcon samples were stored under refrigerated conditions and used for DNA isolation. 1.5 kgs of sterile bag samples were subjected to chemical profiling. Soil chemical profiling involved analysis for pH, electrical conductivity, organic carbon, major nutrients – nitrogen, phosphorus, potassium and minor nutrients – magnesium, calcium, iron, manganese, zinc and copper using standard methods (Singh et al., 2023).

### DNA isolation, quality check and high-throughput sequencing

Total 36 soil samples were processed for DNA isolation using DNeasy Power soil kit following manufacturer’s protocol. Quality check for the DNA was done using Qubit and the 16S amplicon libraries were prepared using Nextera XT kit (Illumina Inc.). The 16S amplicon library was prepared using 16S V3-V4 region primers. Quality check passed amplicons with Illumina adaptors were then amplified using i5 and i7 primers for adding multiplexing indexes and adaptors according to the standard protocol. Amplicon libraries were then sequenced using Illumina MiSeq platform with 300 x 2 read length.

### *In silico* analysis

For all the 36 samples quality check was done using FastQC (v.0.12.0) and adaptor sequences were removed using cutadapt (v.5.1) (Martin, 2011). Denoising, ASV inference, chimera removal and taxonomic assignment were performed using DADA2 pipeline (v.1.36.0) (Callahan et al., 2016) implemented in R. Taxonomic classification was assigned using the SILVA 138 database (Quast et al., 2012). Alpha diversity was calculated based on Observed ASVs, Chao1 estimator and Shannon diversity index using phyloseq package (v.1.52.0) (McMurdie & Holmes, 2013). Non-parametric Kruskal-Wallis tests were applied to assess differences in alpha diversity between groups (habitat and season). To ensure sequencing depth sufficiency and avoid artifacts due to sampling depth, rarefaction curves were generated using subsampling across multiple rarefaction depth, and observed richness was plotted for each sample. Further, for beta diversity analysis, Bray-Curtis dissimilarity matrices were calculated on relative abundance transformed ASV tables (normalized to proportional abundances). Principal Coordinates Analysis (PCoA) ordination was performed to visualize community-level differences in phyloseq. Statistical differences in microbial community composition across habitats and seasons were tested using PERMANOVA (Permutational Multivariate Analysis of Variance. Bacterial functional potential was predicted from 16S rRNA data using PICRUSt2 (Phylogenetic Investigation of Communities by Reconstruction of Unobserved States) (Douglas et al., 2020). Functional profiles were obtained at KEGG ortholog (KO) and pathway levels. Bray-Curtis dissimilarities were calculated on predicted functional profiles, followed by PCoA ordination and PERMANOVA testing to evaluate functional compositional differences across habitats and seasons. Further, DESeq2 (v.1.48.1) was used to get the differentially abundant KOs (log2FoldChange) amongst niche and seasonal comparisons along with their normalized counts (baseMean), p-value and Wald statistics (stat). KOs with significant differential abundance (|log2FoldChange| > 1; padj < 0.05; baseMean > 50 and |stat| > 2) were then filtered.

## Acknowledgement

The study was supported by a grant obtained from the Ignite Life Science Foundation provided by R. H. Sapat Foundation. We are grateful to both R. H. Sapat Foundation and Ignite Life Science for giving us the opportunity to study a Miyawaki forest ecosystem and its dynamics. We are grateful to Dr. Amit Singh’s lab, Indian Institute for Science, Bangalore for allowing us to use their high-performance computing facility which was extremely significant for the bioinformatics data compilation and analysis. We are thankful to MC Kiran and Rohith Ranjan, from the GIS lab at IIHS for helping us with developing the geo-spatial map of the Miyawaki forest and the nearby locations.

## Author contribution

KB analysed the bacterial metagenomic datasets, consolidated the data, and drafted the manuscript. IS was the overall mentor and coordinator for the soil-specific research and manuscript development. VV conducted soil sampling, performed DNA isolation for sequencing, compiled soil nutritional data, and contributed to manuscript writing and figure developments. RR performed all soil nutritional analyses. AJ was instrumental in getting the Miyawaki forest set up and provided information on the soil management practices used for nurturing the Miyawaki forest. JK wrote the proposal and received the grant, designed the overarching research plan and commented on results and interpretations at various stages.

## Supplementary data

**Supplementary table 1:** Tree species planted in the Nashik Miyawaki forest in present study.

**Supplementary table 2:** Differentially abundant KO IDs in comparisons of FD_FW, FD_GD, FW_GW and GD_GW; F: forest, G: grassland, W: wet season, D: dry season.

Table 1. Nutrient analysis of soil samples.

Table 2. Metadata of soil samples used in the present study

